# Evolutionary dynamics of circular RNAs in primates

**DOI:** 10.1101/2021.05.01.442284

**Authors:** Gabriela Santos-Rodriguez, Irina Voineagu, Robert J Weatheritt

## Abstract

Many primate genes produce non-coding circular RNAs (circRNAs). However, the extent of circRNA conservation between closely related species remains unclear. By comparing tissue-specific transcriptomes across over 70 million years of primate evolution, we identify that within 3 million years circRNA expression profiles diverged such that they are more related to species identity than organ type. However, our analysis also revealed a subset of circRNAs with conserved neural expression across tens of millions of years of evolution. These circRNAs are defined by an extended downstream intron that has shown dramatic lengthening during evolution due to the insertion of novel retrotransposons. Our work provides comparative analyses of the mechanisms promoting circRNAs to generate increased transcriptomic complexity in primates.

## Introduction

An important question in biology is how has the complexity of biological systems expanded while the number of protein-coding genes has remained mostly stable. Through decades of research, it has been shown that increased biological complexity has arisen in part by the dynamic generation of unique cell-specific transcriptomes, and as a consequence of the highly versatile programs of gene expression (***Brawand et al., 2011; Cardoso-Moreira et al., 2019***). However, studies of tissues across distant animal lineages have shown that gene expression is highly conserved between the same tissues in different species (***Brawand et al., 2011; Barbosa-Morais et al., 2012; Merkin et al., 2012; Reyes et al., 2013; Cardoso-Moreira et al., 2019***). Hence, gene expression alone is unlikely to explain the heterogeneous expansion in complexity (as defined by the number of cell types) across vertebrate evolution. Instead, it is becoming increasingly evident that the plethora of post-transcriptional mechanisms (***Gueroussov et al., 2017; Ha et al., 2018; Mattick, 2018; Fiszbein et al., 2019; Cheetham et al., 2020; Ha et al., 2021***) capable of greatly expanding transcriptomic diversity also underlies these advances.

Among these, an intriguing class produced by pre-mRNA processing are circular RNAs (circRNAs) (***Memczak et al., 2013; Zhang et al., 2013; Li et al., 2018b; Gokool et al., 2020b***). These non-coding RNAs can regulate protein localization (***Liu et al., 2019***), miRNA functionality (***Piwecka et al., 2017***) and a range of other processes (***Li et al., 2018b; Gokool et al., 2020b***) enabling increased regulatory complexity, especially in the immune and nervous systems (***Piwecka et al., 2017; Gokool et al., 2020a; Guo et al., 2020***). CircRNAs form by back-splicing whereby an exon’s 3’-splice site is ligated to an upstream 5’-splice site forming a closed circular non-coding RNA transcript (***Barrett et al., 2015; Starke et al., 2015***). Back-splicing occurs co-transcriptionally and is facilitated by inverted repeat elements that promote complementarity between adjacent introns favouring circRNA formation over linear splicing (***Jeck et al., 2013; Liang and Wilusz, 2014; Ivanov et al., 2015***). These RNA-RNA interactions can be facilitated by RNA-binding proteins, such as Quaking (***Conn et al., 2015***), that help stabilize the hair-pin structure promoting circRNA formation.

The production of circRNAs can also arise due to the perturbed expression of trans-factors and the inhibition of the core splicing machinery (***Aktaş et al., 2017; Liang et al., 2017***). These spuriously produced circRNAs are maintained as their circular shape protects them from the activity of cellular exonucleases (***Gokool et al., 2020b***). In contrast, the variable usage of cis-regulatory elements in exons and flanking introns can be selected to promote circRNA expression in a cell-type, condition- or species-specific manner (***Nilsen and Graveley, 2010; Irimia and Blencowe, 2012***). Changes in circRNA expression may therefore represent a major source of species- and lineage-specific differences or error-prone mis-splicing. To provide insight into this quandary, here we describe a genome-wide analysis of circRNAs across physiologically equivalent organs from primate species spanning 70 million years of evolution. Our analysis uncovers extensive evidence species-specific circRNAs that display no evidence of conservation even across relatively short evolutionary time-periods. However, we also identify a small subset of circRNAs that are conserved across tens of millions of years displaying increased inclusion rates across evolutionary time. Our analysis reveals that these circRNAs are flanked by newly inserted transposons that correlate with circRNA genesis and extend intron downstream of circRNA. Overall, our results identify evidence of circRNA conservation within closely related species and identify a reoccurring mechanism that correlates with circRNA genesis facilitating the expansion of transcriptomic complexity of primate cells.

## Results

### A core subset of circRNAs show conserved expression signatures but most are species-specific

To address the outstanding questions about the conservation and functional importance of circRNAs, we collected transcriptomic (RNA-seq) data (***Pipes et al., 2013***) from across 9 tissues from 8 primate species, consisting of 3 old-world monkeys, 2 hominoids, 2 new-world monkeys, and one prosimian (**Supplementary Table 1**). These species were chosen on the basis of the quality of their genomes and their close evolutionary relationships enabling the evaluation of transcriptome changes between species ranging from <3 million years to > 70 million years (see **Figure 1A**). For each species, we considered all primate-conserved internal exons as potential origins of back-spliced junctions with no restrictions on backward exon combination. RNA-seq reads were mapped to exon-exon junctions (EEJs) to determine “percent spliced-in” (PSI) for all circRNA with respect to the linear transcript. We also calculated PSI values for linear splicing of each internal exon and transcript per million (TPM) values to estimate gene expression. Orthology relationships between genes and exons were established to enable direct cross-species comparisons.

**Figure 1.**
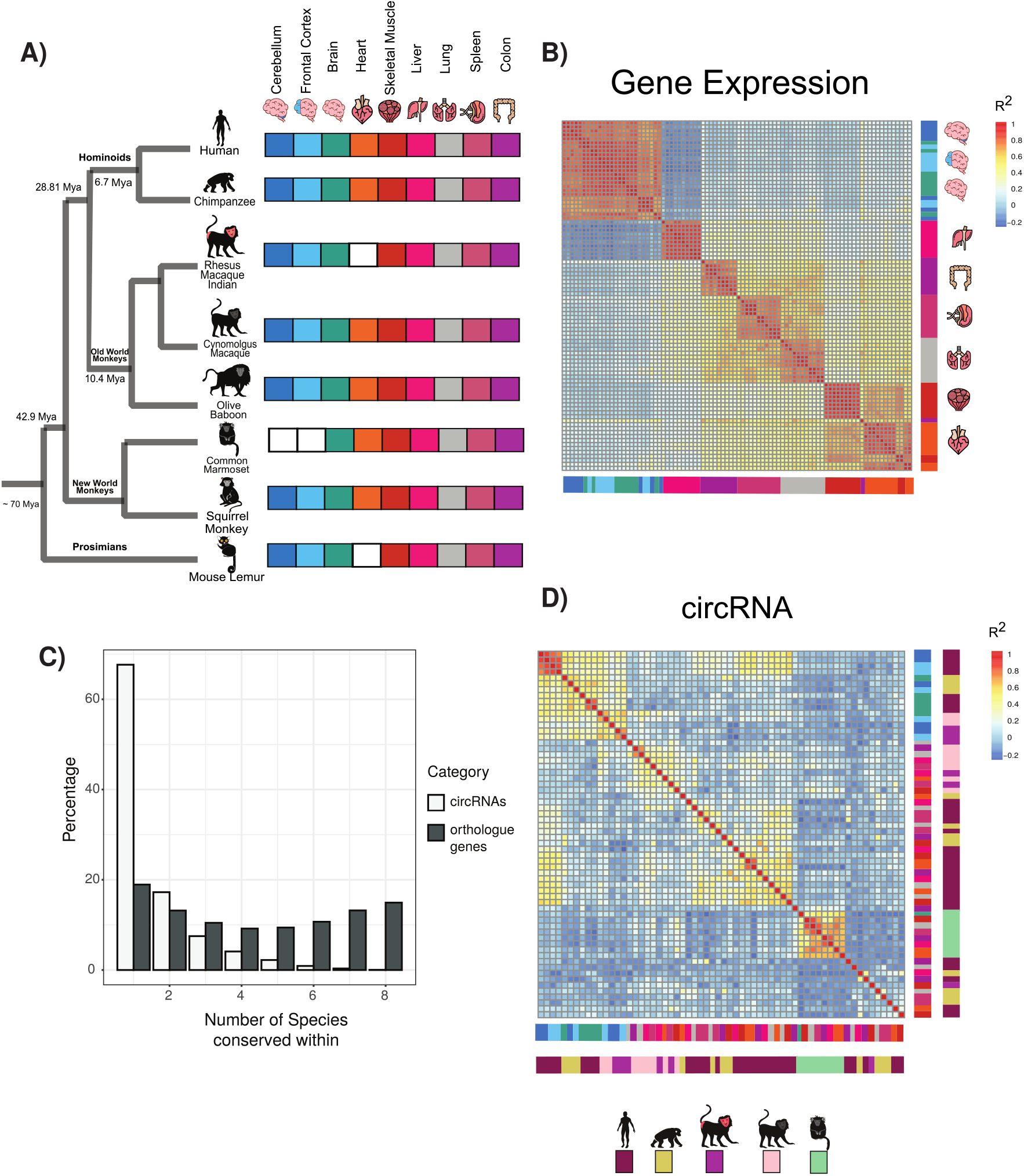
Circular RNA expression signatures are conserved in some tissues. (A) Phylogenetic tree of analyzed species with distance from human in millions of years (Mya) (Divergence time according to TimeTree http://www.timetree.org/). Tissue datasets used in analysis identified on right with white squares denoting lack of dataset. (B) Clustering of samples based on expression values (transcripts per million). The variance of expression values was calculated, and the top 1000 most variable genes were used to calculate the Pearson correlation. (n= 1000 genes in 88 samples). Red colours indicate high correlation between samples and blue describes low correlation. Vertical and horizontal adjacent heatmaps describes tissues (see Fig. 1A for key) (C) Bar plot showing conservation of circRNAs based on back-spliced junction and based on occurrence within orthologous genes.(D) Clustering of conserved circRNAs based on percent spliced in (PSI) values. Clustered using Pearson correlation as in (B). (n=149). Vertical and horizontal adjacent heatmaps describes tissues (inner heatmap(see Figure 1A for key)) and species (outer heatmap).

To initially explore the expression relationships within our datasets we used hierarchical clustering and Pearson’s correlations to determine the gene expression relationships between orthologous genes (see Methods). In agreement with previous results (***Brawand et al., 2011; Barbosa-Morais et al., 2012; Merkin et al., 2012; Reyes et al., 2013***) from analysis across vertebrate species, a clear pattern emerged of tissue-specific conservation of gene expression (**Figure 1B**). This pattern suggests that most tissues possess a tissue-specific gene expression signature such that for example a liver-specific gene in chimp will likely also be liver-specific in lemur. In contrast to previous observations in vertebrates (***Merkin et al., 2012***) there are no clear species-specific exceptions to these patterns likely reflecting the closer evolutionary relationships studied.

To understand circRNA relationships between species, we performed an analogous pairwise clustering analysis using circRNA inclusion values. Replicates from the same tissue invariably clustered together. However, in contrast to gene expression, circRNA expression is segregated by species (**Figure 1-Figure supplement 1A**). This suggests that despite all the exons studied being conserved across primates the majority of circRNAs showed species-specific expression with no orthologous circRNAs in other species (**Figure 1C**, 67% are species-specific, n = 11,201). To evaluate the expression patterns of circRNA orthologs, we identified circRNAs with matched back-spliced junctions (see Methods) conserved across 45 million years of evolution. In this analysis more complex patterns of circRNA conservation emerged with tissue-dominated clustering observed across all types of brain samples (**Figure 1D**). In contrast, for all other tissues circRNAs showed primarily species-specific clustering. Analysis of gene expression changes of genes with these conserved circRNAs (**Figure 1-Figure supplement 1B**,) and alternative splicing changes of exons within conserved circRNAs (**Figure 1-Figure supplement 1C**) showed no consistent changes suggesting circRNA conservation and expression is independent of these regulatory layers.

We next investigated the genes containing circRNAs. Many orthologous genes consistently express circRNAs even if the precise back-spliced junction is not conserved implicating importance of trans-factors in controlling circRNA formation (**Figure 1C**). This phenomenon persisted across species with a median of 10 circRNAs detected per gene across tissues (**Figure 1-Figure supplement 1D**). However, this circRNA production only occurred in a limited number of expressed genes (20.4% of orthologous expressed genes). This suggests certain genomic areas are circRNA factories that are prone to produce large numbers of lowly expressed circRNAs.

These observations suggest a core set of circRNAs show conserved tissue-specific patterns across neural tissues. However, the great prevalence of circRNAs showing species-specific expression indicates that the cis-regulatory or trans-regulatory environments may differ between even very closely related species to promote the species-specific production of circRNAs.

### Features of conserved circRNAs

Our analysis (**Figure 2A**) reveals clear subsets of several hundred circRNAs exhibiting highly conserved circRNA expression. The circRNA ERC1 and many other examples from our data (**Figure 2-Figure supplement 1A, Supplementary Table 2**) demonstrate that circRNA expression can be conserved for tens of millions of years.

To assess the phylogenetic distribution of circRNA across primates we grouped them by PSI values requiring PSI ≥ 5 and at least 5 read support. Out of the approximately 56,000 internal exons with clear orthologs across primates, we identified a large set of circRNA expressing a “species-specific” expression, as well as a set of 773 “conserved circRNAs” that shared expression across at least human, chimp and baboon (**Figure 2-Figure supplement 1B and 1C**). Using our transcriptomic data, we found that a circRNA identified in human was 5-times more likely to be identified in baboon than in lemur, in line with the closer phylogenetic relationship of human to baboon than human to lemur.

**Figure 2.**
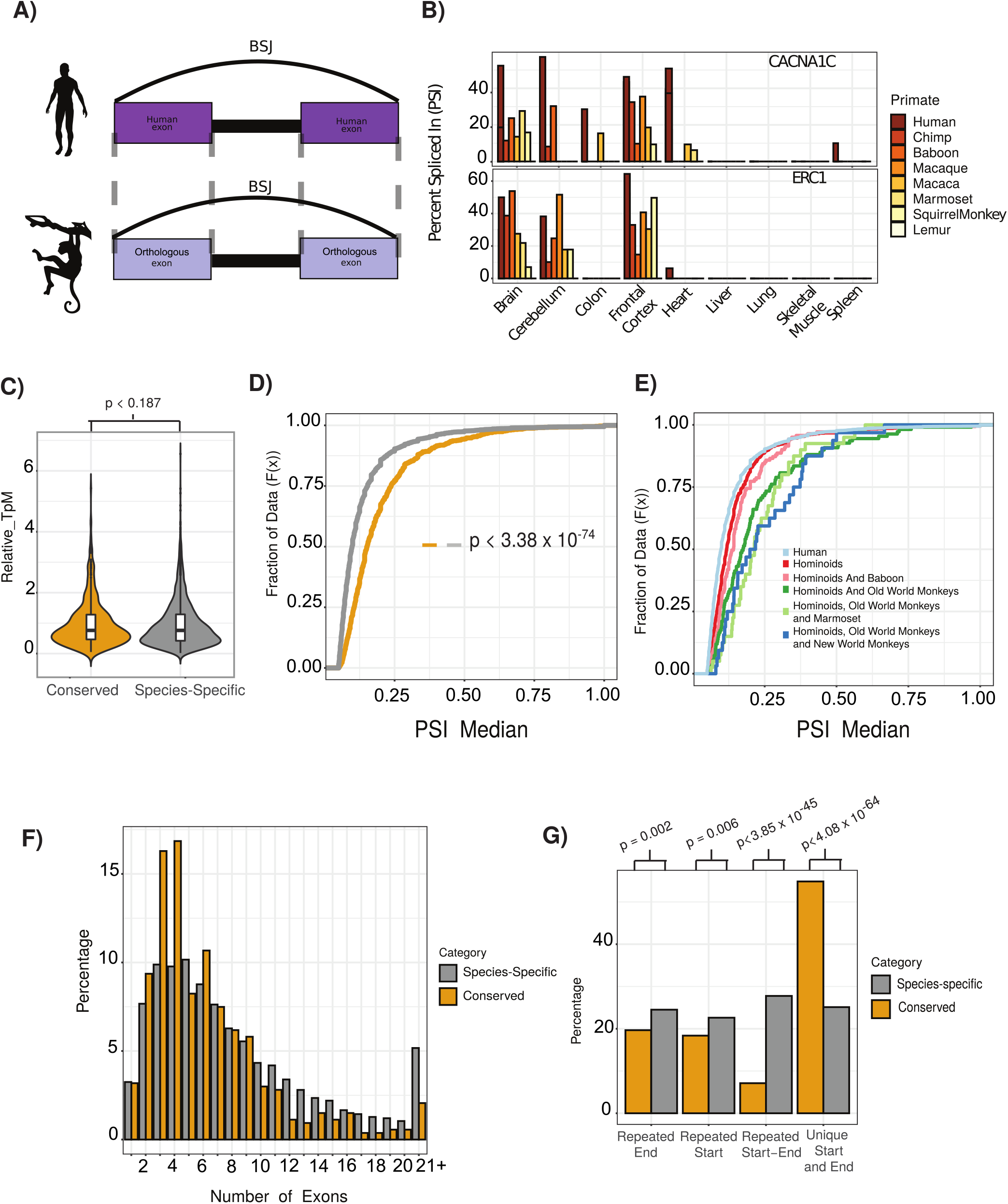
Features of conserved circRNAs. (A) Schematic overview of identification of back-spliced junctions between species. BSJ = back spliced junction. (B) Percent spliced in (PSI) values for conserved circRNAs (top) CACNA1C_chr12:2504436-2512984 and (bottom) ERC1_chr12:1180540-1204512 across tissues and species analyzed. PSI values only calculated for circRNAs with more than 5 reads support. Gene name is indicated in top right-hand corner. (C) Violin plot describing relative expression levels conserved and species-specific circRNAs. Axes on violin plots are Relative TpM values, the normalized circRNA expression according to circRNA reads and gene expression (reads and TpMs values) (see methods). Violin plots show probability densities of the data with internal boxplot. The boxplot display the interquartile range as a solid box, 1.5 times the interquartile range as vertical thin lines and the median as a horizontal line. P-value calculated using Wilcoxon rank sum test (*p <* 0.187). TpM = transcripts per million (D) Cumulative distribution plot of change in percent spliced in (PSI) values across all conserved (yellow) and species-specific (grey) circRNAs. A cumulative distribution plot describes the proportion of data (y-axis) less than or equal to a specified value (x-axis). Cumulative Distribution F(x), cumulative distribution function. p value calculated using Wilcoxon-rank sum test (*p <* 3.38 × 10^−74^). PSI = percent spliced in (E) Cumulative distribution plots of circRNAs with different levels of conservation, as defined by consistent observation of back-spliced junction across species indicated. See 2D for description of cumulative distribution plot. PSI = percent spliced in (F) Bar plot describing number of exons per circRNA for conserved and species-specific circRNAs. Exons are defined by Ensembl and must show evidence of expression (PSI>5 and >5 reads support) in tissue analysed. (G) Bar plot describing uniqueness of start (5’ splice site) and end (3’ splice site) for conserved and species-specific circRNAs. P-values calculated from Fisher’s exact test (*p <* 4.08 × 10^−64^; unique start and end – also see Figure2-Figure supplement 3).

Initial analysis of conserved circRNAs revealed enrichment within neural tissues with over 70% showing consistent tissue expression across 30 million years of evolution (**Supplementary Table 2**). Analysis of expression levels revealed no clear trends for increased expression of conserved circRNAs (**Figure 2-Figure supplement 2A**, *p <* 0.187, Wilcoxon rank sum test vs species-specific), however these circRNAs did display increased inclusion rates, or increased circRNA expression as compared to linear isoform (**Figure 2-Figure supplement 2B**, *p* = 3.38 × 10^−74^ Wilcoxon rank sum test vs species-specific). Furthermore, this inclusion (or circularization) increased with the conservation age of the circRNA (**Figure 2E**, *p* = 8.07 × 10^−19^ Wilcoxon rank sum test of Hominoids vs species-specific (Human specific); *p* = 2.14×10^−06^ Wilcoxon rank sum test of Hominoids vs shared until New World Monkeys). This suggests over time these circRNAs are increasingly influencing the transcriptomic abundance of the linear isoform and the protein abundance of the gene.

Anaylsis of the exonic structure of conserved circRNAs, showed that conserved circRNAs contain fewer exons (**Figure 2F**, *p* = 2.23 × 10^−20^ Wilcoxon rank sum test), and rarely overlap with other circRNAs (**Figure 2G**, *p* = 4.08 × 10^−64^, Fisher exact test; see Methods) displaying back-splicing at unique 5’- and 3’-splice sites. This indicates that these conserved circRNAs possess unique cis- or trans-regulatory features that enable a tight control of the number of exons within a circRNA and the back-spliced junctions used.

### Conserved circRNAs have extensive downstream introns and are flanked by inverted repeat elements

To investigate the role of cis-regulatory elements within conserved circRNAs, we analyzed almost 150 features associated with circRNA formation including a multitude of trans- and cis-regulatory factors and all major groups of transposons (see Methods and **Supplementary Table 3**). To evaluate the influence of these features on defining conserved circRNAs we used two background datasets (see **Supplementary Table 2** and Methods). The first is a background set of randomly combined alternative (10 *< PSI <* 90) exons extracted from genes containing conserved circRNAs (background set). The second is the group of “species-specific circRNAs” defined previously.

Using logistic regression combined with a genetic algorithm for model selection (see Methods), we initially sought to determine the relative contribution of this diverse range of features in defining conserved circRNAs. After initially training our model on a subset of conserved and background circRNAs (80%), we next assessed its performance on the rest of 20% cirRNAs and observed a high average true positive rate of 86.7% (AUC, area under the receiver operating characterstic (ROC) curve) (**Figure 3-Supplementary Figure 1A**) for a model including 24 variables selected by feature analysis. This indicates a core set of 24 cis- and trans-regulatory features drive the conserved formation of circRNAs compared to our background set of introns (**Figure 3A and 3B**). We next used the same approach to determine drivers of conserved and species-specific circRNAs. As expected, our model distinguished these categories less efficiently but was still able to achieve a true positive rate of 65.4% (**Figure 3-Figure supplement 1B**) driven by 12 features. Notable among these features was the depletion of nucleosomes in the downstream intron of the circRNA (**Figure 3-Figure supplement 1D** 1.57 × 10^−03^, Bonferroni-corrected Wilcoxon rank sum test (BH-Wilcox) vs species-specific) and the presence of a more defined 3’-splice site at the final exon (2.04 × 10^−03^, BH-Wilcox vs species-specific).

**Figure 3.**
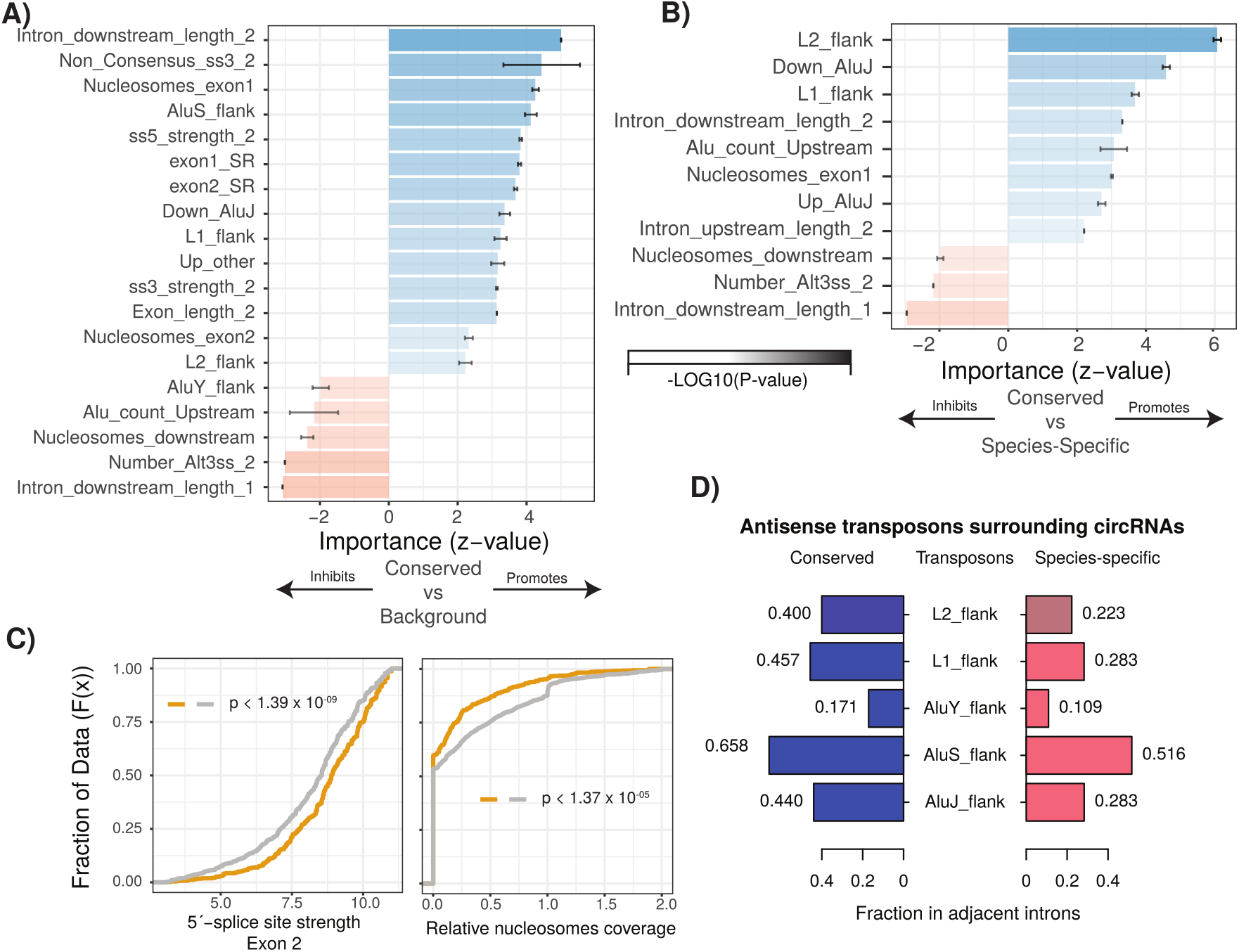
Characterization of cis and trans regulatory features of conserved circRNAs. (A) Bar plot describing feature importance for logistic regression model of conserved circRNAs compared to background. Colours represent positive or negative influence. Transparency reflects log10(p-value of z-statistic). Errors bar represent standard error. “_1” is relative to first exon of circRNA and “_2” is relative to final exon of circRNA. ss3 = 3’ splice site; ss5 = 5’ splice site; Alt3ss = alternative 3’ splice sites. “Flank” are inverted repeats in introns adjacent to circRNAs. See Supplementary Table 3 for details of features. (B) Bar plot describing feature importance for logistic regression model of conserved circRNAs compared to species-specific circRNAs. See 3A for plot interpretation and descriptions. (C) Cumulative distribution plots describing (left; *p <* 1.39*x*10^−09^) 5’ splice site strength at final exon of circRNAs and (right; *p <* 1.37*x*10^−05^) distribution of nucleosomes on intron downstream of circRNA. p-values calculated by Wilcoxon rank sum test and corrected for multi-testing (Bonferroni). See Figure 2D for interpretation of cumulative distribution plot. (D) Pyramid plot showing the mean fraction of circRNAs with selected invertedrepeat retrotransposon elements in adjacent introns.

Introns adjacent to conserved circRNAs also exhibited a significant enrichment for repeat elements (**Figure 3D**, all *p <* 1 × 10^−5^, BH-Wilcox) vs species-specific) in particular L1 and AluJ retro-transposons (**Figure 3D**, L1: *p <* 1.22 × 10^−23^| AluJ: *p <* 1.48 × 10^−18^, BH-Wilcox). A further key distinguishing feature of interest was intron length. Conserved circRNAs exhibited shorter introns downstream of the first exon and an extended intron downstream of the final exon (**Figure 4A and 4B**). In species-specific circRNA this adjacent downstream intron has a median length of 4,624 nucleotides whilst in conserved circRNA the median is almost twice as long at 9,923 nucleotides (**Figure 4B**, *p <* 1.07 × 10^−35^, BH-Wilcox). Finally, when comparing the major drivers of both models, we noticed over 90% (11/12) of features overlapped between the models. This suggests conserved circRNAs are an extreme continuum of species-specific circRNAs. Therefore understanding the processes contributing to circRNA conservation may also provide insight into the genesis of circRNAs across species.

**Figure 4.**
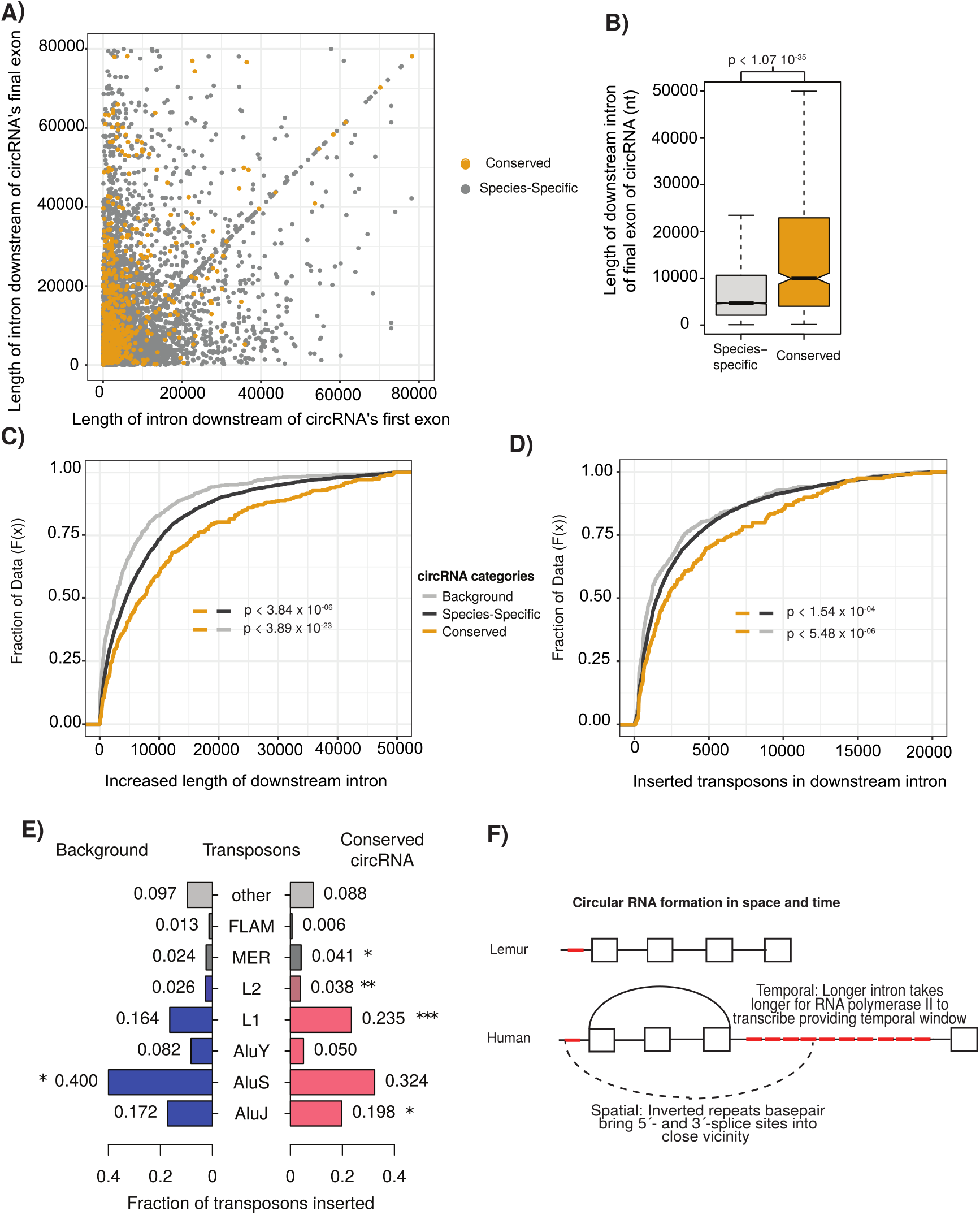
Conserved circRNA downstream intron expanded during primate evolution. A) Scatterplot of downstream intron length for conserved and species-specific circRNAs. (B) Boxplot describing lengths of intron immediately downstream of circRNA for conserved and species-specific circRNAs (see Figure 2C for description of boxplots). p-values calculated by Wilcoxon rank sum test and corrected for multi-testing (Bonferroni). nt = nucleotide (C) Cumulative distribution plot of change of length of orthologous downstream introns of conserved, species-specific and background circRNAs from lemur to human (see Fig. 2D for description of cumulative distribution plots). p-values calculated by Wilcoxon rank sum test and corrected for multi-testing (Bonferroni). (D) Cumulative distribution plot of length of novel repeat elements within the orthologous downstream introns of conserved, species-specific and background circRNAs from lemur to human (see Fig. 2D for description of cumulative distribution plots). p-values calculated by Wilcoxon rank sum test and corrected for multi-testing (Bonferroni). (E) Pyramid plot of the proportion of repeat elements inserted into the downstream introns of conserved, species-specific and background circRNAs from lemur to human. * *−*p <* 0.05; ****−*p <* 0.005, *****−*p <* 1*x*10^−5^. p-values calculated by Wilcoxon rank sum test and corrected for multi-testing (Bonferroni). (F) A schematic model of the results describing impact of our observations on circRNA formation. Boxes represent exons, straight lines are introns, repeat elements are red, arced lines represent back-spliced junction, dashed lines represents RNA-RNA duplex.

### Insertion of young transposons increases downstream intron length in conserved circRNAs

To investigate the evolutionary origins of the switch of conserved circRNAs from absence in prosimians and new world monkeys to conservation within hominoids and old-world monkeys, we investigated the changes in intronic length for the orthologous introns between human (hominoids) and lemur (prosimians). In contrast to orthologous lemur introns, the human introns downstream of all identified circRNAs shows an almost four-fold expansion compared to background dataset of introns within circRNA containing genes (**Figure 4C**, *p <* 3.84 × 10^−23^ Wilcoxon rank sum). This difference is even greater in conserved circRNA, which display an almost 2-fold greater lengthening than species-specific circRNAs (or 8-fold over background) (**Figure 4C**, *p <* 3.84 × 10^−06^, Wilcoxon rank sum). These observations suggest that the expansion of the intron downstream of the circRNA may increase the proportion of backing splicing events increasing the likelihood of circRNA conservation.

To investigate the drivers of this intronic expansion, we aligned the lemur and human introns to identify regions novel to humans. This analysis revealed the insertion of novel transposons at almost double the frequency in introns associated with conserved circRNAs (**Figure 4D**, *p <* 5.48 × 10^−06^, Wilcoxon rank sum). Further evaluation of the retrotransposons revealed this increase in length is driven by the novel insertion of AluJ and L1 elements (**Figure 4E**, AluJ: *p <* 0.018; L1:*p <* 1.73 × 10^−04^, Wilcoxon rank sum). This retrotransposition is potentially facilitated by the depletion of nucleosome occupancy in these introns compared to other human introns (**Figure 3B**, *p <* 1.15 × 10^−07^, BH-Wilcox). Together this argues for the role of young transposons in creating longer intronic regions, which increases the time for RNA polymerase II to reach next canonical splice site and therefore increases likelihood of back-junction splicing to occur.

## Discussion

The evolution of circRNAs has been previously studied across extensive evolutionary time revealing poor conservation for the majority of circRNAs (***Rybak-Wolf et al., 2015; Venø et al., 2015***). Our approach is unique as it focuses on the conservation of circRNAs in very closely related species enabling us to account for the rapid evolution of non-coding RNAs. This increased resolution allowed us to reveal two disparate facts about circRNA expression. Firstly, we observe extensive variation in the production of the vast majority of circRNAs between species. With circRNAs often expressed within the same orthologous genes even if back-spliced junction is not conserved. Conversely, we identify a core set of over 700 circRNA that are conserved across millions of years of evolution. These circRNAs have higher inclusion rates and show increased inclusion across evolutionary age. Both groups are related in the cis- and trans-regulatory features that likely drive circRNA formation such as evidence of recent transposons insertion and extended adjacent introns. However, the conserved groups show decreased diversity of circRNA production and increased expression potentially suggesting a combination circRNA selection and retrotransposon suppression is occurring.

A host of endogenous mechanisms dampen down the impact of the retrotransposons within gene bodies. For example, the formation of Alu exons is suppressed by the nuclear ribonucleo-protein HNRNPC (***Zarnack et al., 2013***) and the nuclear helicase DHX9 binds to inverted repeat Alu elements to suppress circRNA formation (***Aktaş et al., 2017***). Over time though, in selected examples, these inclusions can promote novel functionality (***Shen et al., 2011; Attig et al., 2016, 2018; Avgan et al., 2019***) enabling the creation of tissue-specific exons (***Attig et al., 2018***), miRNAs (***Gu et al., 2009; Spengler et al., 2014***) and promoter regions (***Li et al., 2018a; Zhang et al., 2019***). Our results suggest circRNAs are undergoing a similar selection race with the recent insertion of multiple retrotransposons promoting increased circRNA production that in some cases stabilizes over time. It is important to note though that the production of a large number of circRNAs in itself can be functional (***Liu et al., 2019***). For example, in the immune system a wide diversity of circRNAs are produced and sequester specific RNA binding proteins. These proteins are released upon viral infection to inhibit translation of viral RNA (***Liu et al., 2019***). A major challenge for the field in the following years will arise from determining the contribution of noise versus function for each of these groups.

The investigation of mechanisms controlling circRNA production is a rapid and expanding field (***Li et al., 2018a***). Our results support a kinetic model (***Schor and AR, 2013***) for circRNA function whereby trans-factors promote spliceosome recruitment to the final exon and the very long down-stream introns extend the time-window for back-splicing to occur, which is facilitated by inverted repeats increasing the proximity of 3’-splice site with the upstream 5’-splice site (see Fig. 4F). The extension of the final intron therefore increases the likelihood of circRNA formation in time and space. Spatially by the introducing new retrotransposons, which facilitate RNA-RNA duplex formation (***Jeck et al., 2013; Liang and Wilusz, 2014; Ivanov et al., 2015***) to orientate the splice sites in close proximity, and temporary by increasing the time-window for such an event to occur. The conservation of circRNAs we observe could therefore just be a result of increasing the probability for such an event to occur rather than evidence of functionality. However, circRNAs represent an extreme example of a trend in post-transcriptional regulation whereby low leaky expression creates a pool of possible novel substrates (***Barbosa-Morais et al., 2012; Merkin et al., 2012; Reyes et al., 2013; Mattick, 2018; Avgan et al., 2019; Fiszbein et al., 2019***) increasing the likelihood for unique functionality to arise (***Gueroussov et al., 2017; Guo et al., 2020***). For circRNAs this can be aided by single nucleotide changes that enable trans-acting factors, such as Quaking to facilitate circRNA formation (***Conn et al., 2015***).

In conclusion, our evolutionary analysis identifies that the noisy production of circRNAs is driven by the insertion of novel transposons in adjacent downstream introns that can over time stabilizes to produce conserved circRNAs. This provides a pool of evolutionary potential that could contribute to the evolutionary rewiring of the cell.

## Methods and Materials

### Data processing

All fastq files were quality checked using FastQC (***Andrews, 2010***). Adapters and low quality sequences were removed using Cutadapt (***Martin, 2011***).

### Datasets

Ribo-minus RNA-seq data was extracted from the publically available Nonhuman Primate Reference Transcriptome Resource (NHPRTR) resource (http://www.nhprtr.org/; (***Peng et al., 2015***)). The analyzed samples were from chimpanzee, rhesus macaque, cynomolgus macaque mauritian, olive baboon, common marmoset, squirrel monkey and mouse lemur to cover the 70 MYA of primate evolution (**Supplementary Table 1**). The primates samples of above species were chosen based on the availability of chain files for LiftOver analysis. Human samples were retrieved from different publically available Ribo-minus datasets searching for the SRA IDs in the circAtlas 2.0 database (http://circatlas.biols.ac.cn/; [(***Wu et al., 2020***)]) (**Supplementary Table 1**). Replicates of certain samples across the different primates data were merged to achieve a higher sequencing depth required for alternative splicing quantification (**Supplementary Table 5**).

### Alternative splicing, back-splice junction, and gene expression quantification

Whippet (***Sterne-Weiler et al., 2018***) was used to analyze the RNA-seq samples to quantify casset exon (CE) events, circRNAs (back-spliced junctions; BSJ) and gene expression. To enable BSJ quantification we used the setting with the –circ parameter when running Whippet-quant https://github.com/timbitz/Whippet.jl.

The splice graphs of all primates used for Whippet quantification were calculated using the genome annotation files for each primate from Ensembl (***Yates et al., 2020***) (**Supplementary Table 6**). The genome annotation files were supplemented with novel exon-exon junctions derived from whole genome alignment of primates samples using STAR (***Dobin et al., 2013***) with the 2-pass setting and outFilterMultimapNmax == 10 parameters. Whippet index command was run with the –bam and –suppress-low-tsl parameters.

Gene expression of orthologue genes was retrieved from the gene.tpm.gz files from Whippet-quant output. The correlations of gene expression of orthologue genes between tissue samples from all primates were calculated using the Pearson correlation. Clustering of correlation values was assessed and visualized with a heatmap using the p.heatmap function in R.

### Identification of expressed circRNAs and cassette-exons

All the BSJ events present in orthologue genes between the species mentioned above were filtered to find conserved circRNAs identified by Whippet. The orthologue list of genes was retrieved from Ensembl using the bioMart R package (***Smedley et al., 2009***). Expressed BSJs were defined according to an expression and percent of spliced in (PSI) cutoff of at least 5 reads and ≥ 5% of PSI respectively. Cassette-exon (CE) events from Whippet output were also filtered, keeping those present in orthologue genes and with PSI ≥ 10

### Conservation analysis of circRNAs

We defined a circRNA as conserved if the exon(s) that formed the BSJ are orthologous to the human exon(s) that also formed the BSJ. To achieve this, the exon coordinates of orthologue genes, of each primate were retrieve from the GTF files downloaded from Ensembl (**Supplementary Table 6**). Then, the exon coordinates from the GTF files were intersected with the CE coordinates from Whippet using bedtools intersect (***Quinlan and Hall, 2010***) with -wa parameter.

Then, the resulted exon coordinates (GTF-CE coordinates) were intersected with the circRNAs coordinates within orthologue genes using bedtools intersect with -loj parameter to find which exons were forming the circRNA. The exon coordinates within the circRNA coordinate of the non-human primates were mapped to human coordinates using the UCSC LiftOver (***Navarro Gonzalez et al., 2021***) to retrieve orthologue exons.

The orthologue exons between primates and human were matched to human exon coordinates within the circRNAs coordinates in human to find conserved circRNAs. We defined if a circRNA was conserved between a primate and human if the exon(s) forming the BSJ of the circRNA were also conserved and if the exon(s) start and end coordinates were <= of 100 nc from the start and end of the BSJ coordinate (**Figure 2-Figure supplement 3** for schematic). We defined as non-conserved circRNAs all the human circRNAs that do not have orthologue exons forming the BSJ of the circRNA with other primates.

### Conserved and tissue conserved circRNAs

The list of orthologous circRNAs was plotted in an UpSet plot to visualize the intersection of circRNAs between primates species. We defined the set of conserved circRNAs as the circRNAs within the intersections between primates species where human, chimpanzee and baboon always appeared.

The correlation of inclusion of conserved and tissue-conserved circRNAs between all samples was calculated using the Pearson correlation. Then correlation values were plotted in a heatmap using the p.heatmap function in R.

### Differential gene expression analysis and enrichment analysis of genes with conserved circRNAs

EdgeR (***Robinson et al., 2010***) library was used to perform the differential gene expression analysis between neuronal samples (brain, cerebellum and frontal cortex) and non-neuronal samples (heart, skeletal muscle, liver, lung, spleen and colon). This analysis showed 8,817 differential expressed genes according to a log Fold Change cutoff of log2(1.5) and FDR of 0.05. There were 212 genes of the conserved circRNAs (total of 442 genes) in the set of differential expressed genes. The enrichment of genes with conserved circRNAs was statistically tested with a hypergeometric test using the phyper function in R. The parameters were q = 212, m = 8,817, n = 11,278, k = 442, and lower.tail = FALSE.

### Conserved cassette exons in primates

All exons coordinates of orthologue genes from the GTF files and CE exons coordinates from Whippet were mapped to human coordinates using UCSC LiftOver (***Navarro Gonzalez et al., 2021***). The PSI values of orthologous exons in genes of conserved and tissue-conserved circRNAs were retrieved from all tissues samples of human, chimpanzee and baboon and calculated the Pearson correlation values. The correlation values were plotted in a heatmap using the p.heatmap function.

### Comparison of circRNAs expression and conservation

circRNAs expression of conserved, tissue-conserved and non-conserved circRNAs was calculated using relative TpMs. The relative TpMs were calculated with the below equation.

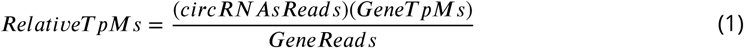

The expression values of conserved and non-conserved circRNAs, and tissue-conserved and non-conserved circRNAs of replicates of the same tissue in human samples were plotted in scatter plots.

The median relative TpM of conserved (and tissue-conserved) and non-conserved circRNAs of human samples were also calculated. The expression values between mentioned sets were statistically compared using a Wilcox test. The parameters of the Wilcox test were x = Conserved (or tissue conserved) circRNAs TpMs, y = Non-conserved circRNAs TpMs, alternative = “greater”. The median relative TpM was plotted in violin plots using the ggplot2 R library (***Wickham, 2016***).

The median PSI values of conserved, tissue-conserved and non-conserved circRNAs across all human samples were calculated. Their inclusion levels were statistically compared using the Wilcox test function in R with the parameters x = Conserved (or tissue conserved) circRNAs median PSI, y = Non-conserved circRNAs median PSI, alternative = “greater”. The distribution of the median PSI values of conserved and non-conserved circRNAs; and tissue-conserved and non-conserved circRNAS were plotted in a cumulative plot using the ggplot2 library in R.

The median PSI value of shared circRNAs between evolutionary interesting sets (human (species-specific circRNAs); hominoids; hominoids and baboon; hominoids and old-world monkyes; hominoids, old-world monkeys and marmoset; and hominoids, old-world monkeys and new-world monkeys) shown in the UpSet plot were calculated, plotted in a cumulative plot and statistically compared using a Wilcox test.

Seven of our reported circRNA from the lists of conserved and tissue conserved circRNAs were of special interest as they were previously reported (***Gokool et al., 2020b***) to be highly expressed in human cerebellum and frontal cortex. The PSI values of such circRNAs were compared across all tissues in the eight primates species.

### Comparison of the number of orthologue genes producing a circRNA and number of conserved circRNAs between species

The number of times an orthologue gene produces at least one circRNA in any of the analyzed species was counted, as well as the number of times a circRNA was shared between another primate. The percentage of shared genes or circRNAs between the eight species was calculated and plotted in a barplot using the ggplot2 library in R.

### Comparison of start and end position of circRNAs between conserved and non-conserved circRNAs

circRNAs can be formed from unique start and end exons forming the BSJ, repeated start exons, repeated end exons, or repeated start and end exons (see **Figure 2-Figure supplement 3** for schematic). The percentage of conserved and non-conserved circRNAs that fall in the above categories was calculated and plotted using the ggplot2 library in R.

### Generalized logistic regression

All continuous data was normalized to ensure a fair comparison between features using scale() package in R environment. Multicollinearity was assessed using the vif() from the R package car. The dataset was split into training (80%) and test (20%). To optimize the selection of the model and the importance of each feature we used the R package glmulti (***Calcagno and De Mazancourt, 2010***). To select from all possible models the selection process used a genetic algorithm (method = ‘g’) with Alkaike information criterion (AIC – crit = “aic”). To calculate the generalized logistic model, glmulti used the R module glm with family = binomial(). ROC curve was calculated using R’s pROC library with test data. Data extracted from this model is reported together with p-value and z-values are reported in **Supplementary Table 7**.

### Background Datasets

Two background datasets were used in this study: background and species-specific (**Supplementary Table 2**). The “background” datasets consisted of exon combinations only within genes with circRNAs. The dataset was constructed by identifying alternative exons within gene of interest (10<PSI<90 within any of the tissues studied) and using python function random to assign these exons together. The “species-specific” dataset was constructed as described above of human circRNA with no evidence of their back-spliced junction being conserved in any other primate species. For both datasets only genes with orthologous genes in all tested primates species were used (based on Ensembl annotation) and only orthologous exons (based on liftover – see above) were used.

### CircRNA features

MaxEntScan (***Yeo and Burge, 2004***) was used to estimate the strength of 3’ and 5’ splice sites. 5’ splice site strength was assessed using a sequence including 3 nt of the exon and 6 nt of the adjacent intron. 3’ splice site strength was assessed using a sequence including - 20 nt of the flanking intron and 3 nt of the exon. SVM-BPfinder (***Corvelo et al., 2010***) was used to estimate branchpoint and polyprimidine tract strength and other statistics. Scores calculated using the sequence of introns to the 3’end of exon between 20 and 500 nt.

Transcription start sites (TSS) were downloaded from Biomart. GC content was calculated using python script. Transposon information download from RepeatMasker as described below. Nucleosome occupancy for HepG2 cells was calculated using data from Enroth et al. (***Enroth et al., 2014***). Colorspace read data was aligned using Bowtie (Langmead, 2010) (-S -C -p 4 -m 3 –best –strata) using index file constructed from Ensembl Hg38. Nuctools (with default settings) was used to calculate occupancy profiles and calculate occupancy at individual regions (***Vainshtein et al., 2017***).

All CLiP-seq data and CHiP-seq data was downloaded pre-processed bed data files from ENCODE (***Sundararaman et al., 2016***) with only narrowpeaks calculated using both isogenic replicates used. Bedtools intersect (-wao) was used to identify overlap with candidate regions. Overlap for all groups of trans-factors were collated and scores normalized by nucleotide length. Groups were based on annotation and split into positive regulators of splicing (SR: Serine/Arginine region containing proteins) and negative regulators of splicing (hnRNP: Heterogeneous nuclear ribonucleoproteins).

In feature analysis, only first and last exons of circRNA, and their surrounding introns, were included in the analysis. The upstream portion is considered as the region 5’ of elements (i.e. first exon) and downstream portion is 3’ of elements.

### Overlap with known repeat elements

Repeat elements identified by RepeatMasker were downloaded from UCSC table browser (***Navarro Gonzalez et al., 2021***) in bed format. Bedtools intersect (-wao) was used to identify overlap of transposons with novel exons.

The frequency of transposable events is calculated as the proportion of transposons overlapping area of interest (i.e. exon 1). All transposons were grouped together into 12 categories (AluJ, AluS, AluY, L1, L2, L3, MIR, MER, FLAM, AT_rich, SINE and everything else into “other”) based on annotation from RepeatMasker. Flanking regions are defined as having the same transposable elements on differents strands in both introns adjacent to the circRNA.

### Intronic length and transposons comparison of human and lemur

Orthologs exons between human and lemur containing circRNAs were identified using the procedure described above. Intron length was determined based on the nearest exon from ENSEMBL annotation (***Yates et al., 2020***) with evidence from RNA-seq data of expression (PSI>10). To identify regions unique to human, the intronic regions unique to human were split into windows of 20 nucleotides. Liftover was used to identify conserved regions between human and lemur genomes for each of these windows. Regions with no evidence of conservation were overlapped (using bedtools intersect –wao) with UCSC RepeatMasker (***Navarro Gonzalez et al., 2021***) annotation to identify novel transposon insertion.

## Supplementary Data

### Supplementary Figures

Supplementary Figures are available as an attachment to this document.

### Supplementary Tables

Supplementary Tables are available as a separate attachment to this manuscript.

**Supplementary Table 1** Primates datasets IDs and sequencing depth information.

**Supplementary Table 2** Conserved, non-conserved (human specific) and background circRNAs.

**Supplementary Table 3** Features associated with circRNA formation (trans- and cisregulatory factors and all major groups of transposons).

**Supplementary Table 4** Primates specific circRNAs.

**Supplementary Table 5** Information about merged samples to acquire higher sequencing depth.

**Supplementary Table 6** Genome version, GTF and chain file information.

**Supplementary Table 7** GLM output.

## Acknowledgments

We gratefully acknowledge John Mattick, Akira Gookol, Juli Wang and Helen King for helpful discussions and feedback on this study, as well as all members of the Weatheritt Lab. G.S.R was supported by a UNSW UIPA PhD Scholarship. This research was supported by the NSW Institute of Cancer Research (RJW), the Scrimshaw Foundation (RJW), the Australian Research Council (ARC) Discovery Project (RJW, IV), an ARC future fellowship (IV) and a University of New South Wales Scientia Fellowship (IV).

## Author contribution

Contributions to this publications are distributed as follows: Study design: G.S.R, I.V., R.W.; Bioinformatic data analyses: G.S.R and R.W.; Paper manuscript and discussion: G.S.R, I.V. and R.W.

## Supplementary Figures

**Figure 1–Figure supplement 1.**
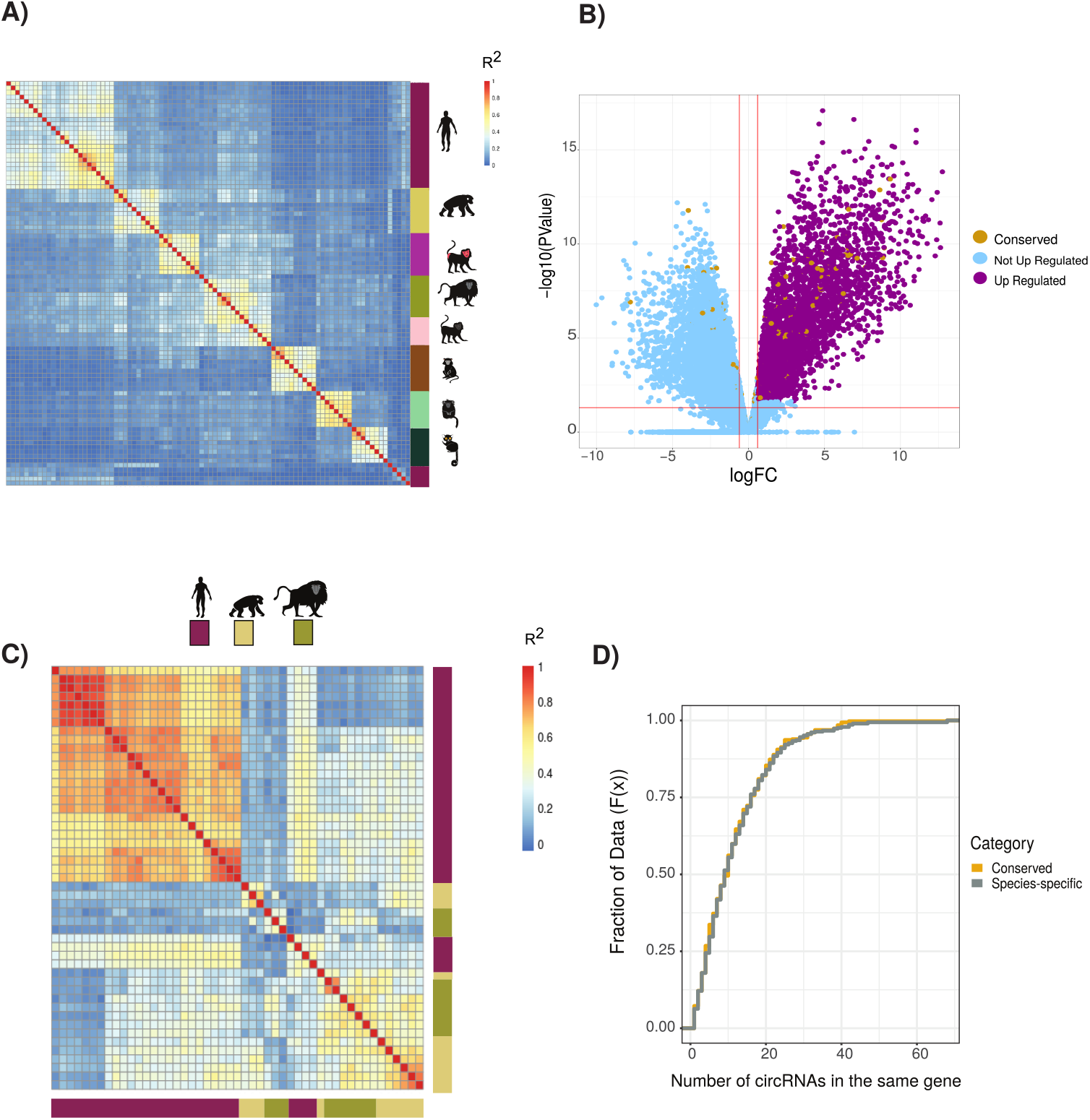
(A) Clustering of circRNAs based on percent spliced in (PSI) values. Clustered using Pearson correlation as in Fig. 1B. (n=19,005). Vertical heatmap indicates primate species. See Fig. 1B for details of heatmap and Supplementary Table S4 for data used. (B) Volcano plot of differential gene expression analysis between neuronal samples and non-neuronal samples. In blue are “not up-regulated genes” (n = 15,661), in purple are “up-regulated genes” (n = 4,434) and in yellow the “genes with conserved circRNAs” (n = 442). Likelihood of circRNA genes being enriched in differentially expression genes (p = 0.036, hypergeometric test). FC = fold value. P-value in figures calculated by quasi-likelihood negative binomial test and corrected for multi-testing (Bonferroni) (C) Clustering of alternative splicing events based on percent spliced in (PSI) of exons within conserved circRNAs shows no clustering by tissue. (n = 1,256). Vertical and horizontal adjacent heatmaps represents species. See Fig. 1B for details of heatmap. (D) Cumulative distribution plot displaying the number of circRNAs found within same gene. (see Figure 2D for description of cumulative distribution plots)

**Figure 2–Figure supplement 1.**
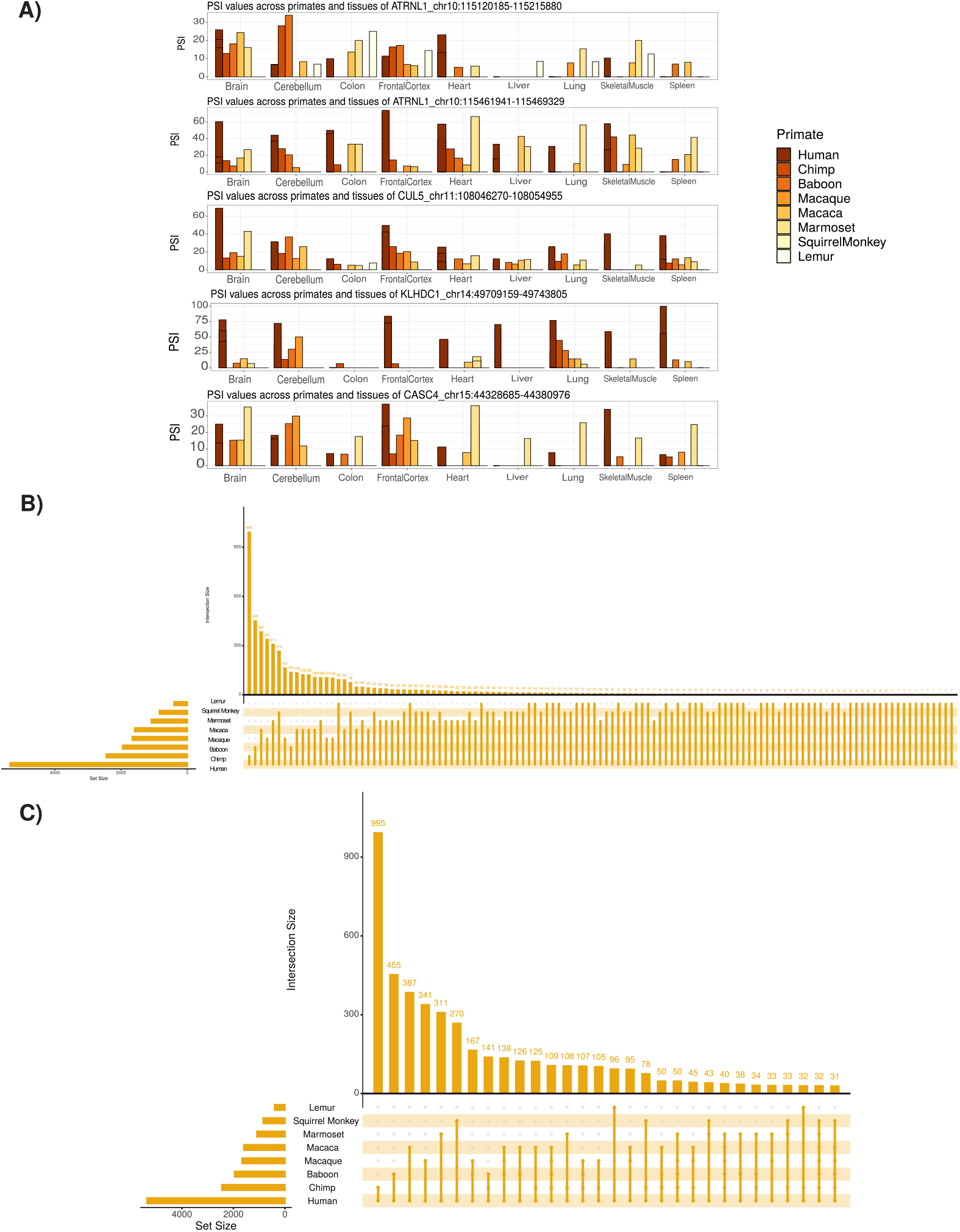
(A) Extension of Figure 2B showing examples of identified circRNAs, Coordinates for each circRNA are shown. PSI = percent spliced in (B) Upset plot of conserved circRNAs across primate species analysed. An upset plot displays the intersections of a set. Each column corresponds to a set, and each row corresponds to one segment in a Venn diagram. Number of top of bars represent number in each overlap. (C) Upset plot of conserved circRNAs across primate species studied with at least 30 overlap. (see Figure 2-Supplementary Figure 1B for description of Upset plot).

**Figure 2–Figure supplement 2.**
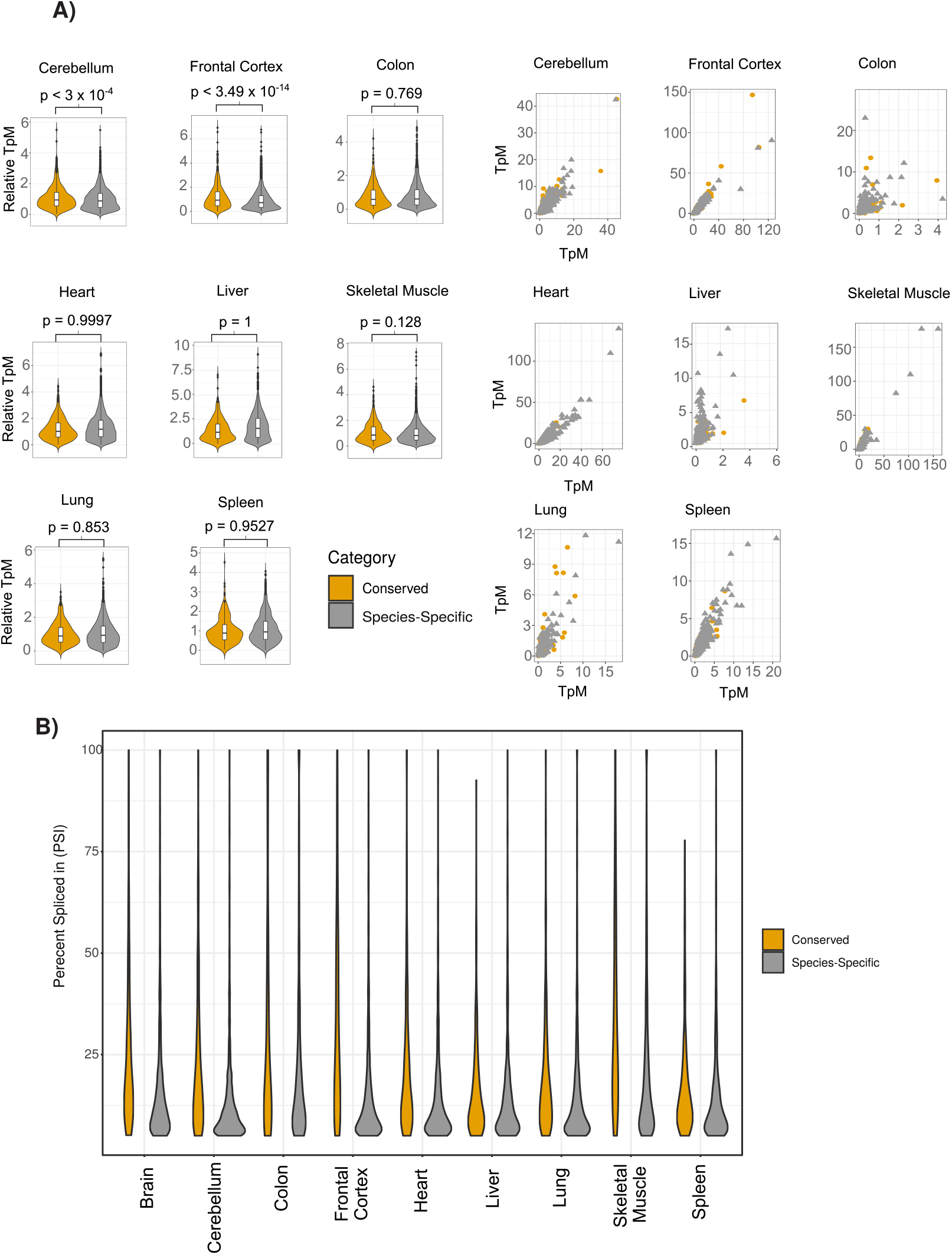
(A) Extension of Figure 2C showing violin plots (left) and scatter plots (right) across all human tissues. Axes on violin plots are Relative TpM values, the normalized circRNA expression according to circRNA reads and gene expression (reads and TpMs values) (see methods). P-value is calculated using Wilcoxon rank sum test. (see Figure 2C for description of violin plots) (B) Extension of Figure 2D showing violin plots of percent spliced in (PSI) value differences across all individual tissues (see Figure 2C for description of violin plots)

**Figure 2–Figure supplement 3.**
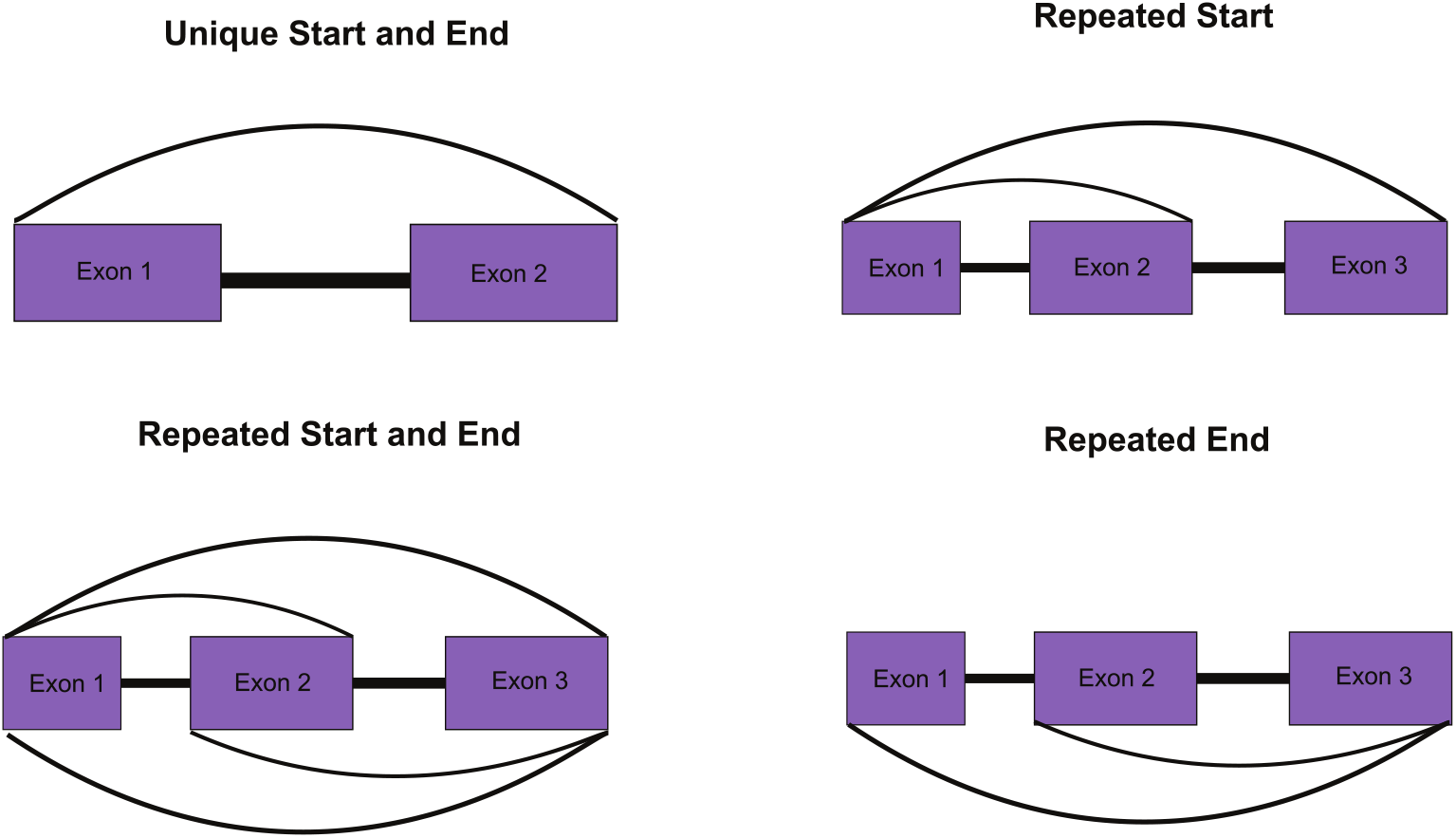
Overview of approach to identifying unique circRNAs for Figure 2G (see Methods for details)

**Figure 3–Figure supplement 1.**
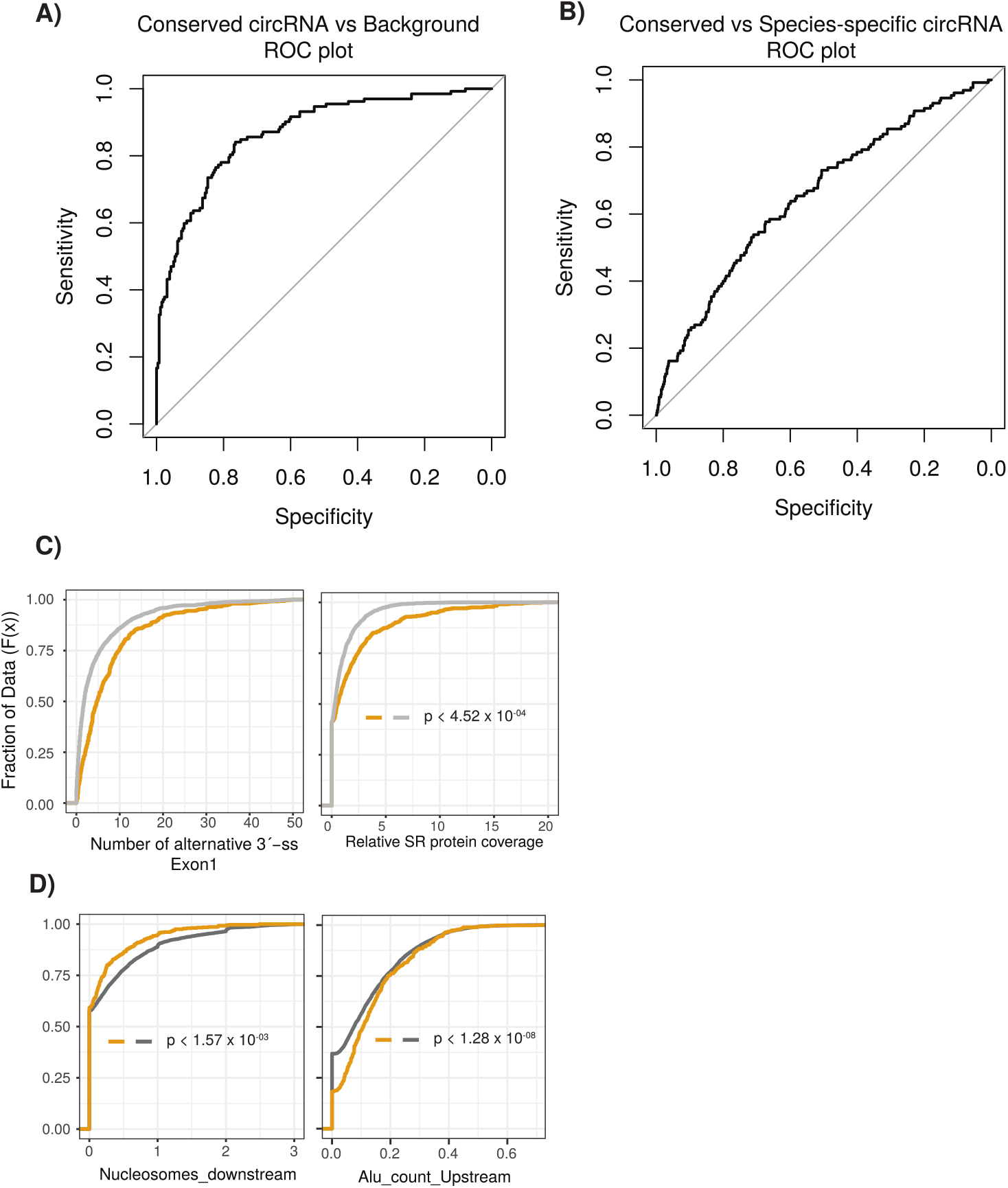
(A) An ROC (Receiving operating characteristic curve) plot displaying the sensitivity (true positive rate) compared to the selectivity (false positive rate) for the logistic regression model (conserved versus background circRNA). (B) An ROC plot for logistic regression model (conserved versus speciesspecific circRNA). (C) Cumulative distribution plots comparing conserved versus background circRNAs of (left) alternative 3’ splice sites (ss) within first exon of the circRNA and (right) of Serine/Arginine (SR) RNA-binding peaks from CLiP data. p-values calculated by Wilcoxon rank sum test and corrected for multi-testing (Bonferroni). (see Figure 2D for description of cumulative distribution plots). (D) Cumulative distribution plots comparing conserved versus species-specific circRNAs of (left) nucleosome peaks in downstream intron adjacent to circRNA and (right) of Alu element content in upstream intron adjacent to circRNA. (see Figure 2D for description of cumulative distribution plots). p-values calculated by Wilcoxon rank sum test and corrected for multi-testing (Bonferroni).

